# BTS: scalable Bayesian Tissue Score for prioritizing GWAS variants and their functional contexts across omics data

**DOI:** 10.1101/2024.10.30.621077

**Authors:** Pavel P. Kuksa, Matei Ionita, Luke Carter, Jeffrey Cifello, Kaylyn Clark, Otto Valladares, Yuk Yee Leung, Li-San Wang

## Abstract

**Motivation:** Summary statistics from genome-wide association studies (GWAS) are widely used in fine-mapping and colocalization analyses to identify causal variants and their enrichment in functional contexts, such as affected cell types and genomic features. With the expansion of functional genomic (FG) datasets, which now include hundreds of thousands of tracks across various cell and tissue types, it is critical to establish scalable algorithms integrating thousands of diverse FG annotations with GWAS results.

**Results:** We propose BTS (Bayesian Tissue Score), a novel, highly efficient algorithm uniquely designed for 1) identifying affected cell types and functional elements (context-mapping) and 2) fine-mapping potentially causal variants in a context-specific manner using large collections of cell type-specific FG annotation tracks. BTS leverages GWAS summary statistics and annotation-specific Bayesian models to analyze genome-wide annotation tracks, including enhancers, open chromatin, and histone marks. We evaluated BTS on GWAS summary statistics for immune and cardiovascular traits, such as Inflammatory Bowel Disease (IBD), Rheumatoid Arthritis (RA), Systemic Lupus Erythematosus (SLE), and Coronary Artery Disease (CAD). Our results demonstrate that BTS is over *100x* more efficient in estimating functional annotation effects and context-specific variant fine-mapping compared to existing methods. Importantly, this large-scale Bayesian approach prioritizes both known and novel annotations, cell types, genomic regions, and variants and provides valuable biological insights into the functional contexts of these diseases.

**Availability and implementation:** Docker image is available at https://hub.docker.com/r/wanglab/bts with pre-installed BTS R package (https://bitbucket.org/wanglab-upenn/BTS-R) and BTS GWAS summary statistics analysis pipeline (https://bitbucket.org/wanglab-upenn/bts-pipeline).

## Introduction

Typical single-marker-based genome-wide association studies (GWASs) do not consider correlation between variants (linkage disequilibrium, LD) and the potential cellular or epigenetic context of the genetic variants. Interpretation and prioritization of GWAS results and variants thus require downstream analyses using various types of functional genomic (FG) data.

Many existing methods can perform fine-mapping of the GWAS signals within a region, based on the summary statistics alone (approximate Bayes factor), or summary statistics with LD information. Examples of these methods include conditional analysis (Galarneau et al., 2010; Knight et al., 2012; Uffelmann et al., 2021; Yang et al., 2012), CAVIAR/CAVIARBF (Chen et al., 2015; Chen et al., 2016; Hormozdiari et al., 2014), DAP-G (Wen et al., 2016), FINEMAP (Benner et al., 2016), and SuSiE (Wang et al., 2020; Zou et al., 2022). Other methods prioritize variants based on their functional annotations, such as coding, promoter or enhancer regions, e.g., RegulomeDB (Boyle et al., 2012; Dong et al., 2023). Colocalization methods such as coloc (Giambartolomei et al., 2014; Wallace, 2021), HyPrColoc (Foley et al., 2021), eCaviar (Hormozdiari et al., 2016), ENLOC (Wen et al., 2017) and their variants jointly analyze summary statistics from GWAS and molecular traits such as expression quantitative trail loci (eQTL). There are also tools for integrative analysis (e.g., INFERNO (Amlie-Wolf et al., 2018), SparkINFERNO (Kuksa et al., 2020), FUMA (Watanabe et al., 2017)) which combine data from multiple sources: for example, prioritizing variants based on both colocalization posterior probabilities and overlap with functional annotations. Additionally, some methods (e.g., PAINTOR (Kichaev et al., 2014), fGWAS (Pickrell, 2014), BFGWAS_QUANT (Chen et al., 2022), PolyFun (Weissbrod et al., 2020), CARMA (Yang et al., 2023)) further formalize overlap with functional annotations and incorporate functional annotations as priors for inferring causal variant status.

The recent increase in availability of large functional annotation databases such as ENCODE (Consortium, 2012; Consortium et al., 2020), ROADMAP (Consortium et al., 2020; Roadmap Epigenomics et al., 2015), and FILER (Kuksa et al., 2022) provides an opportunity to carry out unbiased functional analyses of GWAS results across thousands of cell types and genome-wide annotations. However, such large-scale, systematic analyses can come at a significant computational cost, as they need to build and evaluate different statistical models for each annotation or combination of annotations and cell types of interest. In addition, each model evaluation often requires running time exponential in the number of potentially causal variants (Asimit et al., 2019; Kichaev et al., 2014).

Moreover, most methods that model LD are susceptible to errors due to a mismatch between GWAS summary statistics and LD. Such mismatches can occur when the GWAS and the reference genotype panel used to compute LD differ demographically. Even more importantly, mismatch is all but guaranteed in the case of GWAS meta-analyses, as reviewed recently in the SLALOM publication (Kanai et al., 2022). Briefly, if two variants belong to the same haplotype, most models expect them to have similar summary statistics and behave erratically if this expectation is violated. But different studies in a meta-analysis can incorporate one variant and not the other, which can cause large differences in the amount of evidence supporting each variant. To accommodate this common situation, fine-mapping methods must be robust to LD mismatch (Chen et al., 2021; Kanai et al., 2022; Yang et al., 2023).

To address these large-scale analysis issues, we present BTS, a Bayesian Tissue Score model and an efficient implementation of this model. BTS performs joint fine-mapping of variants and their context-mapping based on FG annotations and provides easy-to-interpret summaries of the results. The main features of the proposed BTS framework are:

- BTS can perform joint context-mapping (inference of cell types, genomic features) and context-specific fine-mapping of variants (**Figs. 1,3,5; Methods**)
- End-to-end GWAS summary statistics analysis pipeline (**Fig. 1**; **Section “**BTS GWAS summary statistics analysis workflow“; **Supplementary Methods**): users can provide their own functional annotations for running with BTS, or use the FILER FG database (Kuksa et al., 2022) to obtain cell type-specific annotations from data sources such as ENCODE (Consortium, 2012; Consortium et al., 2020), EPIMAP (Boix et al., 2021), and GTEx (Consortium, 2020). By default, the user only needs to provide GWAS summary statistics as input.
- Scalability: BTS can conduct a systematic and exhaustive search through thousands of genome-wide annotation tracks and has running times two orders-of-magnitude faster per genome-wide track (**Section** “Running time improvement”; **Fig. 6**) using a novel, more efficient factored Bayesian model (**Methods**; **Section** “BTS statistical model”) for FG annotations, LD and GWAS summary statistics.
- Robustness to mismatch between GWAS summary statistics and LD estimates (**Fig. 2**). The BTS model introduces a single model parameter (the prior on variance of the true effect sizes) to control most of the sensitivity to LD mismatch. We also provide guidelines on how to choose this parameter. (**Methods; Section** “Mitigation of LD mismatch”).

**Figure 1.**
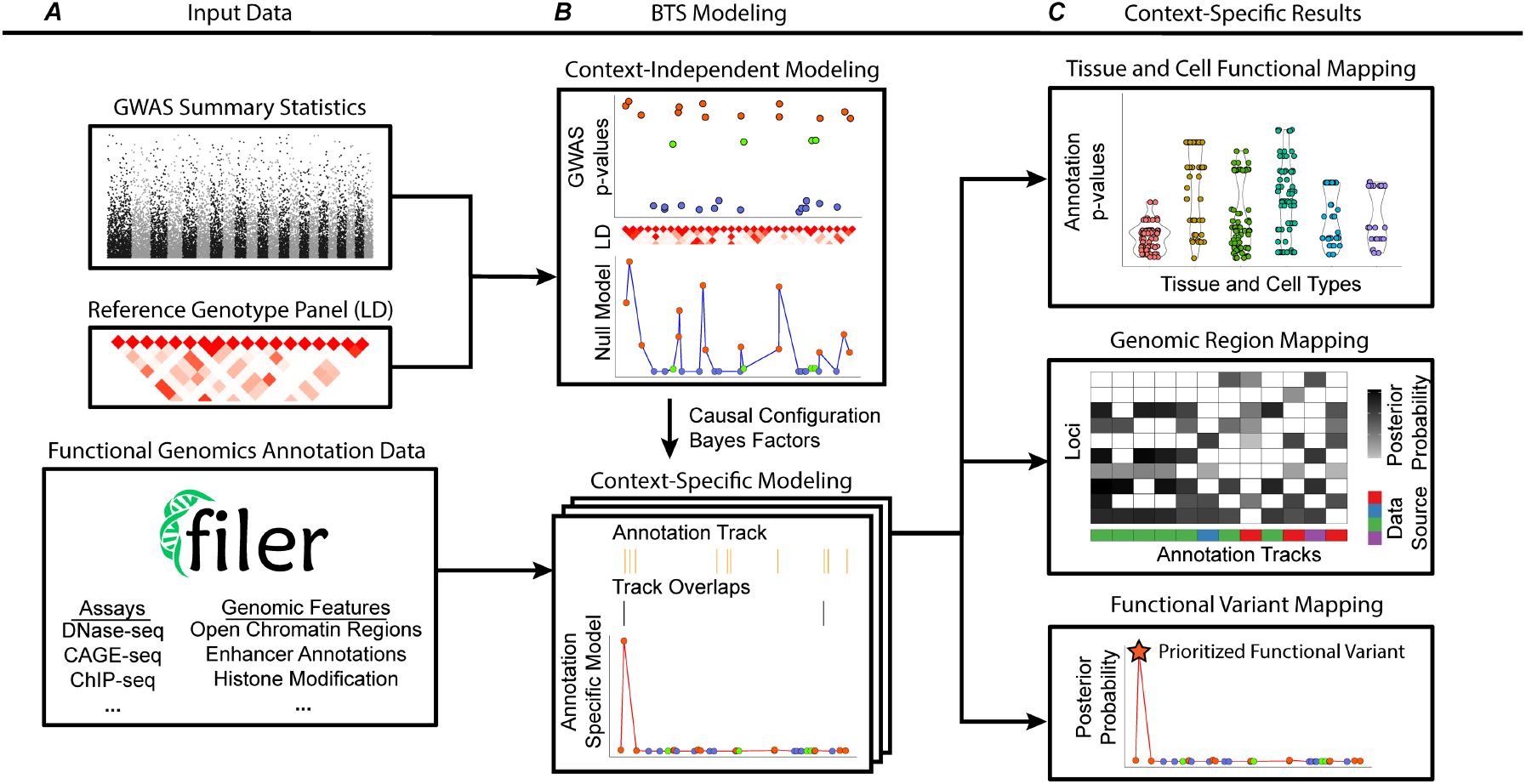
Overview of the BTS framework. **A)** Input GWAS summary statistics and LD data (top sub-panel) are analyzed across cell type and tissue-level FG annotations (bottom sub-panel) to jointly perform context-mapping and identify context-specific set of genomic regions and potentially causal variants for each context and region. **B)** The two-factor structure of BTS model (**Section** “BTS statistical model”) allows it to be evaluated efficiently across thousands of genome-wide annotations (**Fig. 6; Section** “Running time improvement”). BTS estimates context-independent GWAS+LD null model (top sub-panel) and many annotation-specific models (bottom sub-panel). **C)** BTS outputs annotation relevance (enrichment *p*-values) (top sub-panel), functional genomic regions within each of the prioritized contexts (middle sub-panel), and annotation-specific causal variant posterior probabilities (bottom sub-panel) (**Fig. 3**; **Sections “**Prioritizing regions, variants and their contexts with BTS**”, “**Cross-trait BTS **evaluation“**). **Supplementary Fig. S1** and **Section** “BTS GWAS summary statistics analysis workflow” detail BTS analysis workflow and its main steps.

**Figure 2.**
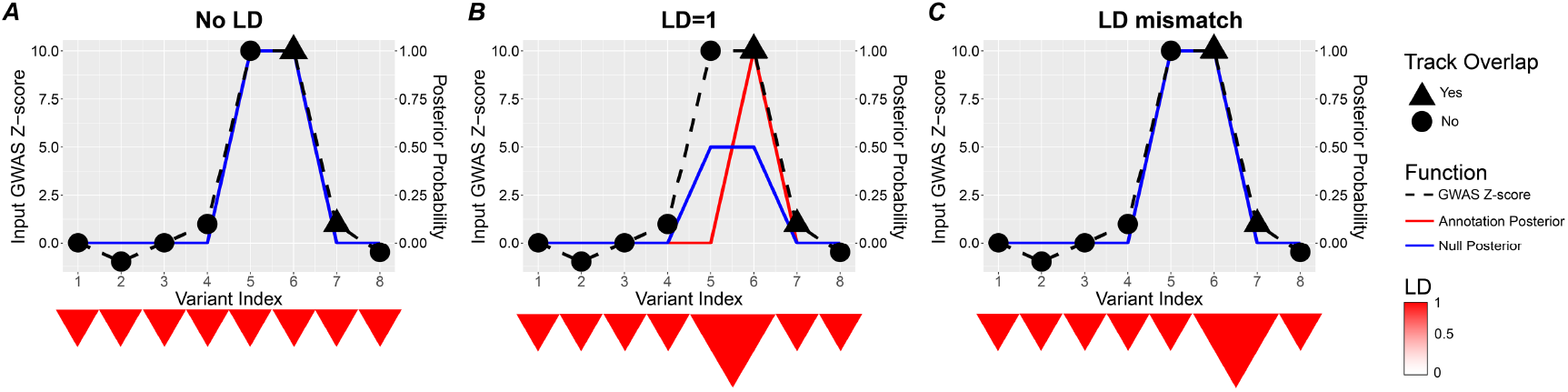
Variant prioritization with BTS using summary statistics, LD, and annotation information. **A**) In case of two independent signals (numbered 5 and 6 on the x-axis), both variants are prioritized by BTS (posterior probability for both variants is 1). **B**) If the two variants (5 and 6) are in LD, variant 6 is prioritized (red line posterior probability) based on its overlap with relevant annotation track (black triangle). Note that the null model (blue line) will not be able to prioritize between these variants (posterior=0.5) since both variants have similar Z-scores and are in LD. **C**) In case of LD and GWAS mismatch, two variants (6 and 7) are in LD but have very different GWAS summary statistics (Z-scores). As shown here, BTS is robust with respect to the LD mismatch and prioritizes variant 6 with a more significant association statistic (Z-score) and avoids prioritizing false-positive variant 7 by default model.

We applied BTS to GWAS datasets from four different diseases: Coronary Artery Disease (CAD) (van der Harst & Verweij, 2018), Inflammatory Bowel Disease (IBD) (Liu et al., 2015), Rheumatoid Arthritis (RA) (Stahl et al., 2010) and Systemic Lupus Erythematosus (SLE) (Bentham et al., 2015). In each case, BTS took under one hour (**Fig. 6**) to prioritize cell types, tissues, genomic regions and variants relevant to the disease across a variety of FG annotation tracks (>900; see **Supplementary Table S2** for details on annotation tracks used). This shows BTS can serve as a tool for systematic and unbiased functional context mapping and context-specific variant mapping.

The BTS GWAS summary statistics analysis pipeline, including scripts for obtaining relevant LD and functional annotations starting from the GWAS summary statistics, is freely available at https://bitbucket.org/wanglab-upenn/BTS-pipeline. BTS model estimation is implemented as an R package and is freely available at https://bitbucket.org/wanglab-upenn/BTS-R. BTS Docker including pre-installed GWAS summary statistics pipeline is also available at https://hub.docker.com/r/wanglab/bts.

## Results

### BTS overview

To jointly estimate variant posterior probabilities and enrichment of annotations in causal variants, we rely on a well-known Bayesian model (PAINTOR (Kichaev et al., 2014), CAVIARBF (Chen et al., 2015), see also **Methods; Section “**BTS statistical model**”; Supplementary Methods**). Briefly, for each possible configuration of causal variants within a locus, its Bayes factor only depends on the GWAS summary statistics and the correlation between variants (LD). On the other hand, causal variant configuration prior depends only on the functional annotations of the variants, as well as the annotation importance (e.g., enrichments of the annotations in causal variants). We use an Expectation-Maximization (EM) algorithm to maximize overall data likelihood by iteratively updating the annotation enrichments and evaluating configuration posterior probabilities, until a convergence criterion is reached (**Fig. 1; Methods; Supplementary Methods**).

Our key observation is that the posterior probability decomposes into a factor which only involves the GWAS data, and one which only involves annotations (**Section “**BTS statistical model**”**; **Eq. 9**). Furthermore, the factor involving GWAS data is the same for all iterations of the EM algorithm. Because of this, BTS can compute Bayes factors only once and then re-use them hundreds or thousands of times across various annotation evaluations (**Figs. 1, 6**).

BTS algorithm further improves running time by introducing two main computational improvements 1) by using the matrix inversion lemma to compute Bayes factors (**Section “**BTS statistical model**”; Supplementary Methods; Lemma 2, Eq. 4**) and 2) by more efficiently computing variant configuration probabilities (**Section “**BTS statistical model**”; Eq. 8**).

These observations, alongside computational improvements detailed in the **Methods, and Supplementary Methods**, are the main reason for the improved runtime of BTS (**Fig. 6**).

To run the BTS algorithm, the user has the option to supply three types of information for each genomic region to be analyzed: per-locus GWAS summary statistics, LD matrix, and functional annotations. Alternatively, the user can provide full GWAS summary statistics only and use BTS end-to-end GWAS summary statistics analysis pipeline (**Fig. 1**) which (1) identifies genomic regions of interest, (2) computes per-locus variant LD and functional annotation matrices, then (3) runs BTS model estimation on the assembled input data (Steps 3,4 in **Supplementary Fig. S1**), and (4) aggregates across estimated models to prioritize annotations, genomic regions, and variants. This GWAS summary statistics analysis pipeline leverages the FILER FG data repository (Kuksa et al., 2022) for tissue and cell type-level functional annotations, and the 1000 Genomes project genotype reference panel (Byrska-Bishop et al., 2022; Genomes Project et al., 2015) to estimate LD between variants. The pipeline outputs cell type and functional context-specific posterior probabilities for individual variants within each of the identified genomic regions of interest, annotation and functional context importance (enrichment scores, causal variant prior odds), as well as cell type, region, and variant mapping summary plots (e.g., **Figs. 3, 4**)

**Figure 3.**
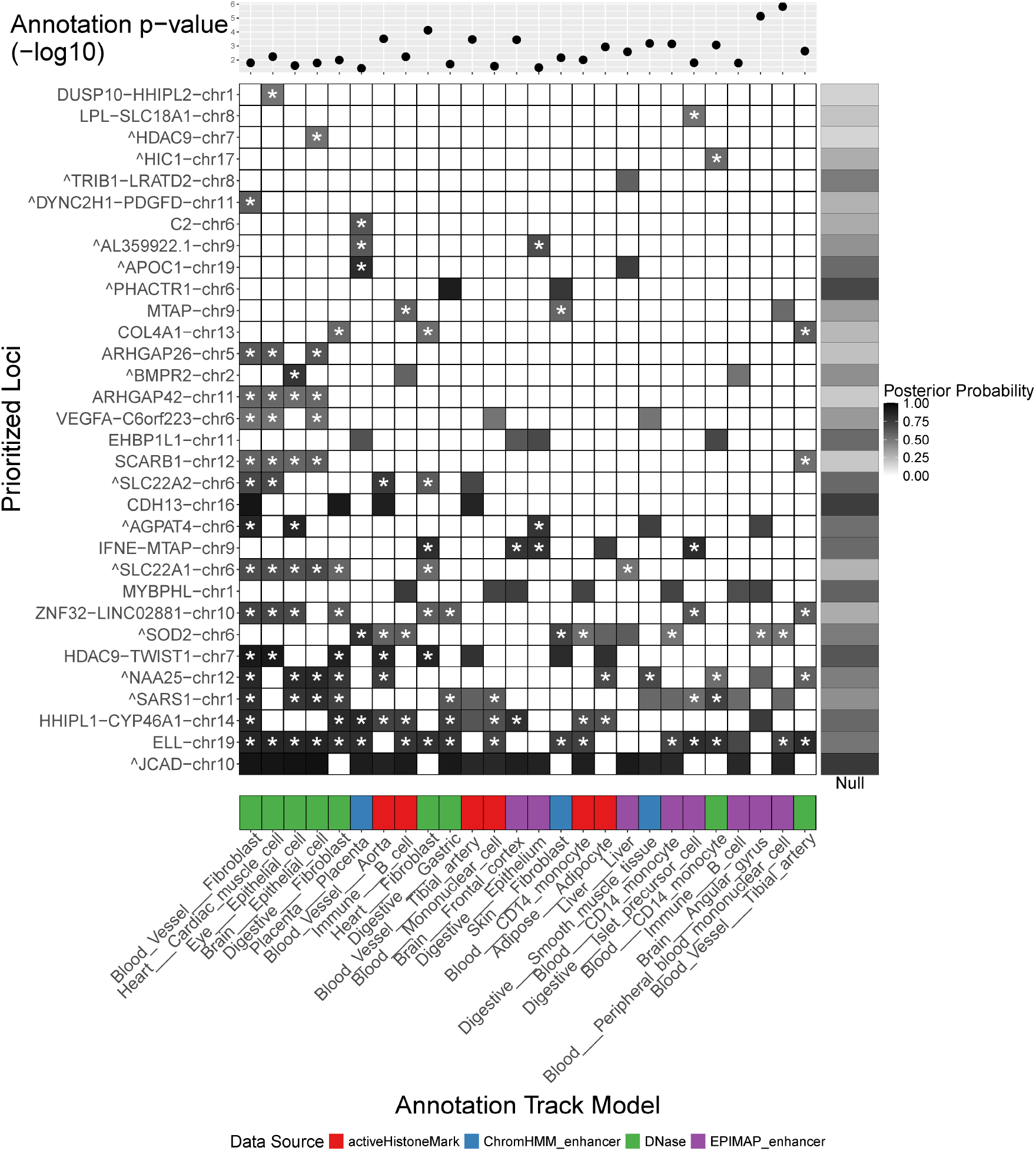
BTS prioritization for functional contexts (X axis) and genomic regions (Y axis) in CAD GWAS. **A**) Effect size and enrichment of the functional annotation genome-wide across all identified regions of interest and using all variants located within these regions. Shown are the annotation significance (*p*-values) as given by the likelihood ratio test of the BTS model with annotation and the model without annotation (see **Section** “BTS algorithm“; **Supplementary Methods** for details on estimating annotation significance). Out of 103 significant annotations (*p* < 0.01), shown are 25 annotations overlapping with at least five prioritized genomic regions (variant posterior > 0.5). In general, annotations with greater enrichment in top GWAS variants have lower, more significant *p*-values. **B**) BTS-prioritized regions of interest (Y axis) and their potential functional contexts (X axis). Shown are regions with annotation overlaps and top variant posterior > 0.5. Asterisks (*) mark regions with increased causal variant posterior compared to the null model (GWAS+LD only, without annotation) in one or more functional contexts. Darker colors correspond to regions and annotations with greater top variant posteriors. Merged regions (i.e. regions containing more than one overlapping LD blocks) are labeled with a caret (^) before the region name. For CAD, BTS prioritizes blood vessel, monocyte, cardiac muscle, and immune cell types across active histone marks, open chromatin, and enhancer genomic features.

**Figure 4.**
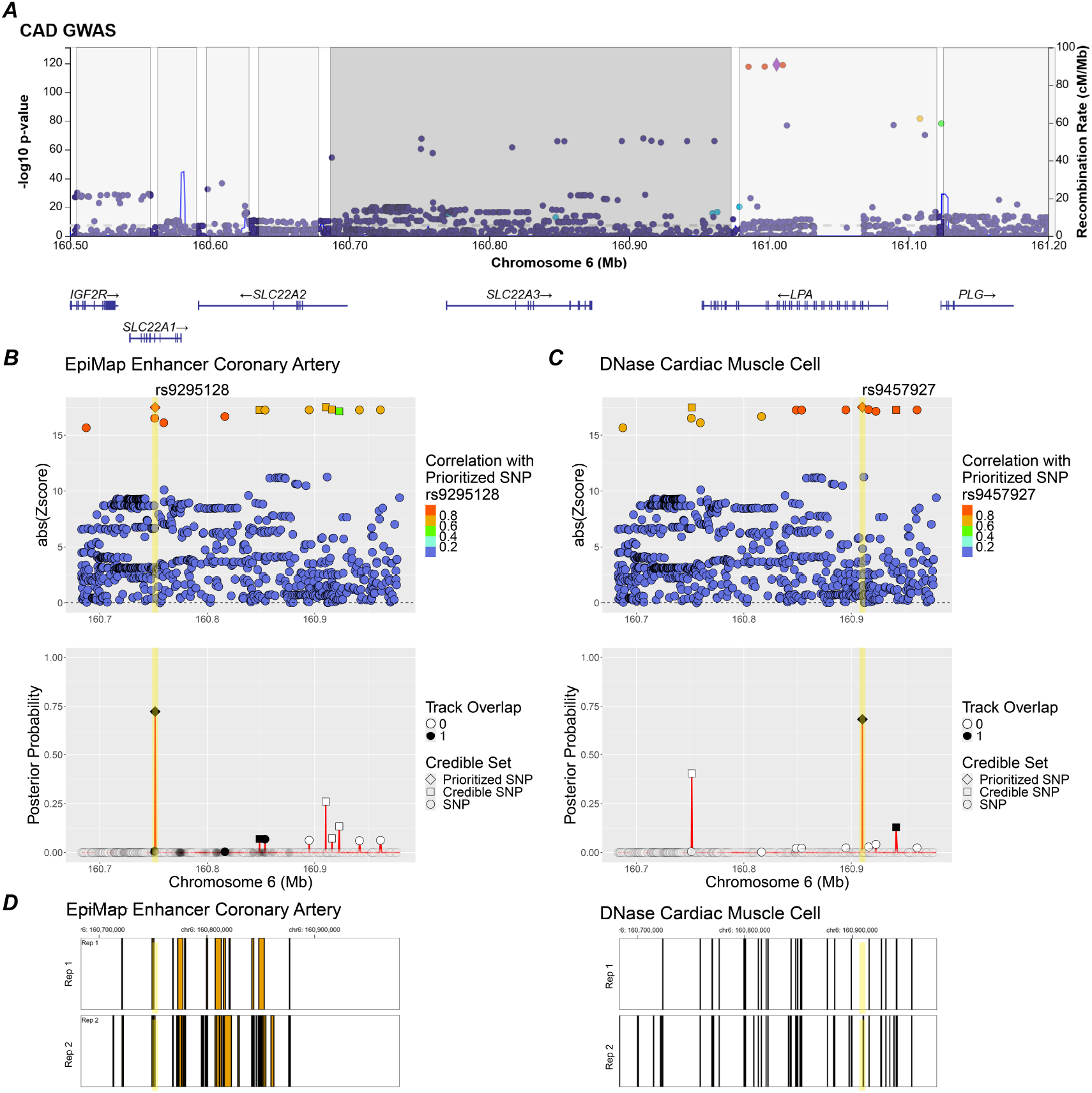
BTS within-region context-specific variant fine-mapping. Shown is an example of differential variant prioritization for SLC22A2 region (GRCh37/hg19 chr6:160683145-160978997) in two different contexts, coronary cell enhancers and open chromatin regions in cardiac muscle cells. **A)** GWAS results for genomic region encompassing SLC22A, with discrete LD blocks shaded separately. Subfigures **B, C, D** show BTS analysis results for the middle (dark grey) LD block. **B)** BTS prioritizes variant rs9295128 in coronary artery context as it overlaps with annotated enhancer from EpiMAP (BTS coronary artery enhancer annotation *p*-value=1.7e-3; prior odds=2.85). **C)** In a different context (heart), BTS prioritizes variant rs9457927 in the cardiac muscle cell based on its location within an annotated open chromatin region defined by DNase assay (BTS annotation *p*-value=7.1e-4; prior odds=5.51). **D)** Annotated enhancer regions for coronary artery (left) and open chromatin regions for cardiac muscle cell (right) across two biological replicates (Rep1, Rep2) each.

As illustrated in the example in **Fig. 2**, given the same summary statistics, BTS discovers two independent signals if the variants are uncorrelated (**Fig. 2**, first column, LD=0), and one independent signal shared between two variants, if these are correlated (**Fig. 2**, second column, LD=1). Crucially, in the baseline model with no functional annotations, the posterior probability mass is shared equally between the two variants, but in the model that uses an annotation the probability is assigned to the variant which overlaps trait-relevant annotation.

Finally, BTS is robust to mismatch between LD and summary statistics (**Fig. 2**, third column, LD mismatch): when two variants with LD close to 1 have wildly different summary statistics, the one with low phenotype association is not prioritized. We emphasize that this common-sense result was difficult to achieve: the same statistical model with different parameter settings has a propensity for prioritizing variants with low phenotype association in such cases of LD mismatch. One of our contributions in this article is to identify parameter values that make the model robust to LD mismatch (**Supplementary Methods; Methods “**Mitigation of LD **mismatch“**).

### Prioritizing regions, variants and their contexts with BTS

To illustrate the performance of BTS, we started with summary statistics from a Coronary Artery Disease (CAD) GWAS (van der Harst & Verweij, 2018). Our end-to-end pipeline first performed LD pruning of the 4,298 genome-wide significant GWAS variants (*p-*value*<5e-8)* to identify 389 tag variants (pairwise-independent, r^2^<0.7). We further filtered out 2 loci belonging to the HLA region on chromosome 6, where complicated linkage patterns make fine-mapping extremely difficult. LD expansion of the remaining tag variants yielded 6,513 candidate variants. Using the LD-expanded candidate set, we then defined the LD block for each tag variant using the left-most and right-most variants in LD with the tag variant 1) within a 1 megabase window and 2) with no more than 1000 variants separating the boundary variants and the tag variant. Any of the overlapping LD blocks were then merged to obtain 167 non-overlapping regions of interest. By extracting variants within these regions from the initial GWAS summary statistics, we obtained a total of 24,727 variants for further analysis. We note that each of the regions will not only contain variants that are highly associated with the CAD phenotype or are in LD with them but also any unrelated or unassociated variants in between. For each of the identified regions of interest, the BTS pipeline used PLINK (Chang et al., 2015; Purcell et al., 2007) to compute LD matrices based on the 1000 Genomes dataset (Byrska-Bishop et al., 2022; Genomes Project et al., 2015), using genotypes from samples belonging to the European super-population as reference.

BTS used FILER (Kuksa et al., 2022) to obtain a discovery set of 943 various regulatory annotations for testing including FANTOM5 enhancers (Andersson et al., 2014), Roadmap Epigenomics ChromHMM enhancers (Roadmap Epigenomics et al., 2015), ENCODE (Consortium, 2012; Consortium et al., 2020) DNase hypersensitivity sites, EpiMap enhancers (Boix et al., 2021) and all available tissues and cell types therein (see **Supplementary Tables S2, S4**). After excluding annotations which did not overlap any of the regions of interest, 532 genome-wide annotation tracks remained, across all data sources and tissues. Then BTS was run on the 167 genomic regions, with the default setting of *d*=2, i.e. looking for at most two distinct causal signals in each region (we note that even though *d*=2, the credible variant set may include more than *d* variants).

BTS prioritized open chromatin annotations for blood vessel, monocyte and B cells and other relevant cell types and tissues (**Fig. 3**) consistent with the tissues and cell types involved in the disease (Ghattas et al., 2013; Libby & Theroux, 2005; Srikakulapu & McNamara, 2017). Blood vessel, blood, and immune-related annotations accounted for 23 out of 103 (22%) of the significant annotations (annotation statistical significance assessed with a likelihood ratio test, *p*<0.01).

Importantly, BTS provides context-specific variant fine-mapping as shown in **Fig. 4**, where BTS prioritized two variants (rs9295128 and rs9457927) within the same SLC22A2 region, with one variant (rs9295128) prioritized in coronary artery context, and another variant (rs9457927) prioritized in the cardiac muscle cell.

Using cell type and tissue-specific annotations with BTS helped identify a more precise set of potentially causal variants across genomic regions for CAD. Compared to the GWAS+LD null model without annotation, we observed a significant reduction in average credible set sizes for BTS models that use annotations. For example, across 46 genomic regions prioritized by BTS and 200 annotations we observed the average credible set size of 2.44±2.7, while the null model had a significantly larger credible set size of 6.5±7.77 across the same genomic regions. Importantly, using annotations allowed BTS to identify 44 additional potentially causal variants that were otherwise not prioritized by the null model. We also note 38 out of 72 variants identified by GWAS+LD-only model were de-prioritized as they were not found to be located within any of the prioritized annotations or functional elements. Overall, using annotations with BTS allowed to identify a total of 78 potentially causal CAD variants across 46 loci (see **Supplementary Table 5a**).

### Cross-trait BTS evaluation

We further tested performance of BTS on several other GWAS datasets for immune-related traits (IBD, RA, SLE) using the same set of 943 annotation tracks as input. As shown in **Fig. 5**, BTS prioritizes disease and cell type associations consistent with previous studies, including myeloid dendritic cells for IBD (Baumgart et al., 2005), T cells for RA (Weyand et al., 2000), and monocytes for SLE (Hirose et al., 2019) (ChromHMM enhancers) and CAD (Ghattas et al., 2013) (active histone marks, enhancer and DNase-hypersensitive regions). See **Supplementary Tables 5a-d** for detailed list of all prioritized annotations, genomic regions, and variants for each tested GWAS dataset.

**Figure 5.**
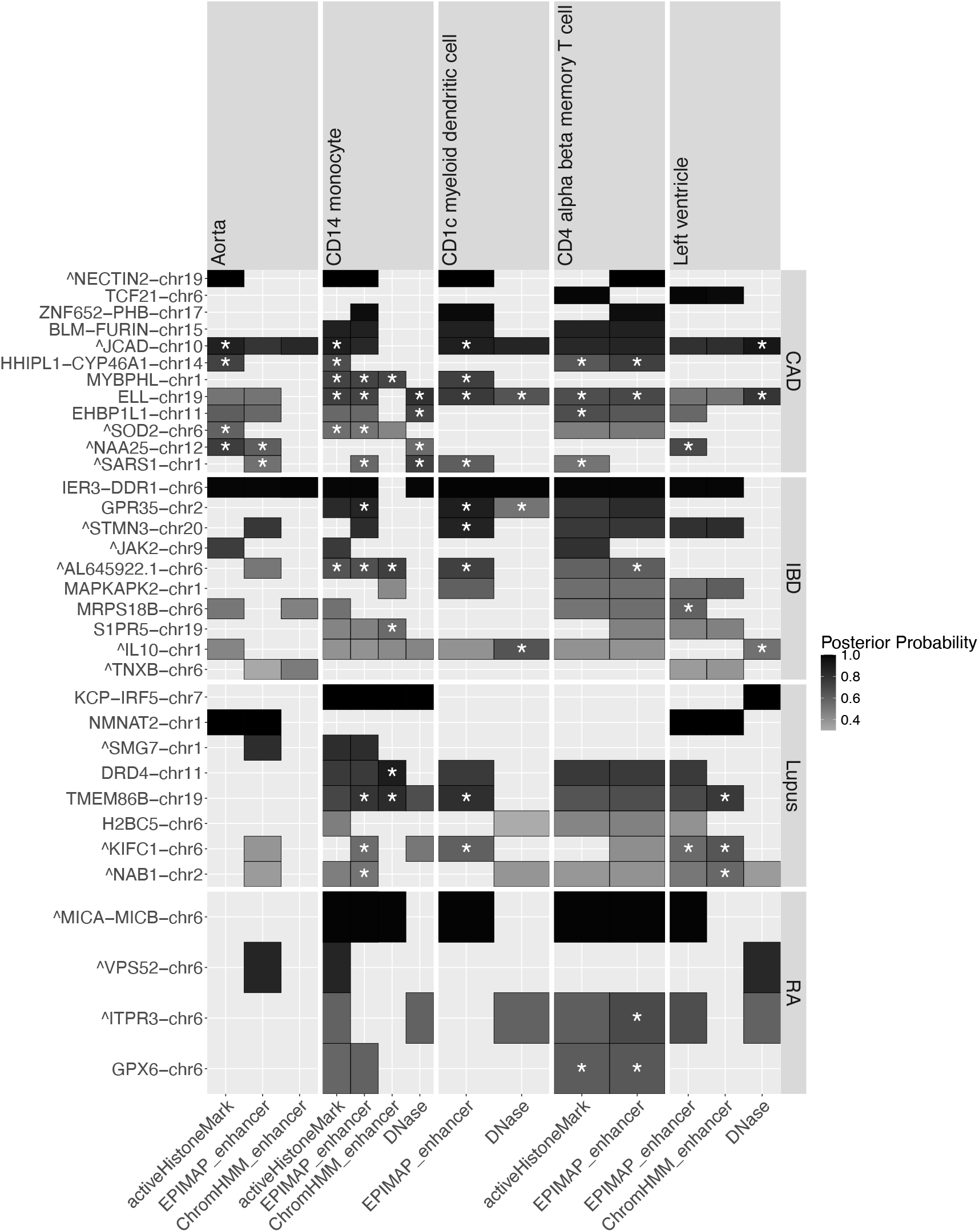
BTS results across four immune-related and cardiovascular GWAS traits. Shown are identified functional contexts (X axis) and genomic regions (Y axis) at the cell-type level for each of the traits (CAD, IBD, SLE, and RA panels on the right). For each region and functional context, the top variant posterior is shown (shades of gray) with a star (*) indicating posterior increase of at least 0.2 in that context compared to the null model without annotation. BTS identifies specific cell types and genomic feature types (open chromatin, histone marks, enhancers on the X axis) for each of the traits (e.g., CD4 T cell for RA; myeloid dendritic cells for IBD; aorta blood vessel and CD4 alpha-beta T cells for CAD; heart left ventricle for SLE).

Moreover, BTS prioritized regions which have been implicated in IBD (**Supplementary Fig. S2**). Among these are an intergenic region near the gene coding for the transcription factor CEBPA (Zhou et al., 2019), or genes coding for interleukins and their receptors, such as IL10 (Ip et al., 2017) and ILR1 (Dosh et al., 2019). Importantly, while recent functional work on IBD (Stankey et al., 2024) identified intergenic region directing macrophage inflammation through ETS2, the ETS2-PSMG1 region was prioritized by BTS in granulocytes (**Supplementary Fig. S2**), belonging to the same myeloid family with macrophage.

### Running time improvement

BTS only took minutes to process genome-wide GWAS data, which involved tens of thousands of variants and hundreds of annotation tracks (**Fig. 6**). Compared to reference implementation fastPaintor/Paintor v3.0 (Kichaev et al., 2017), BTS is faster by two orders of magnitude (**Fig. 6c; Section** “BTS statistical model”), being able to compute an annotation-specific model in under one second on average (**Fig. 6a**,**b**). This comes at the cost of a pre-processing step that takes a few seconds but is only carried out once for the entire analysis. Moreover, BTS scales linearly (**Fig. 6a**,**b**) with both the number of genomic regions and the number of annotation tracks, which guarantees reasonable running times for even larger datasets.

**Figure 6.**
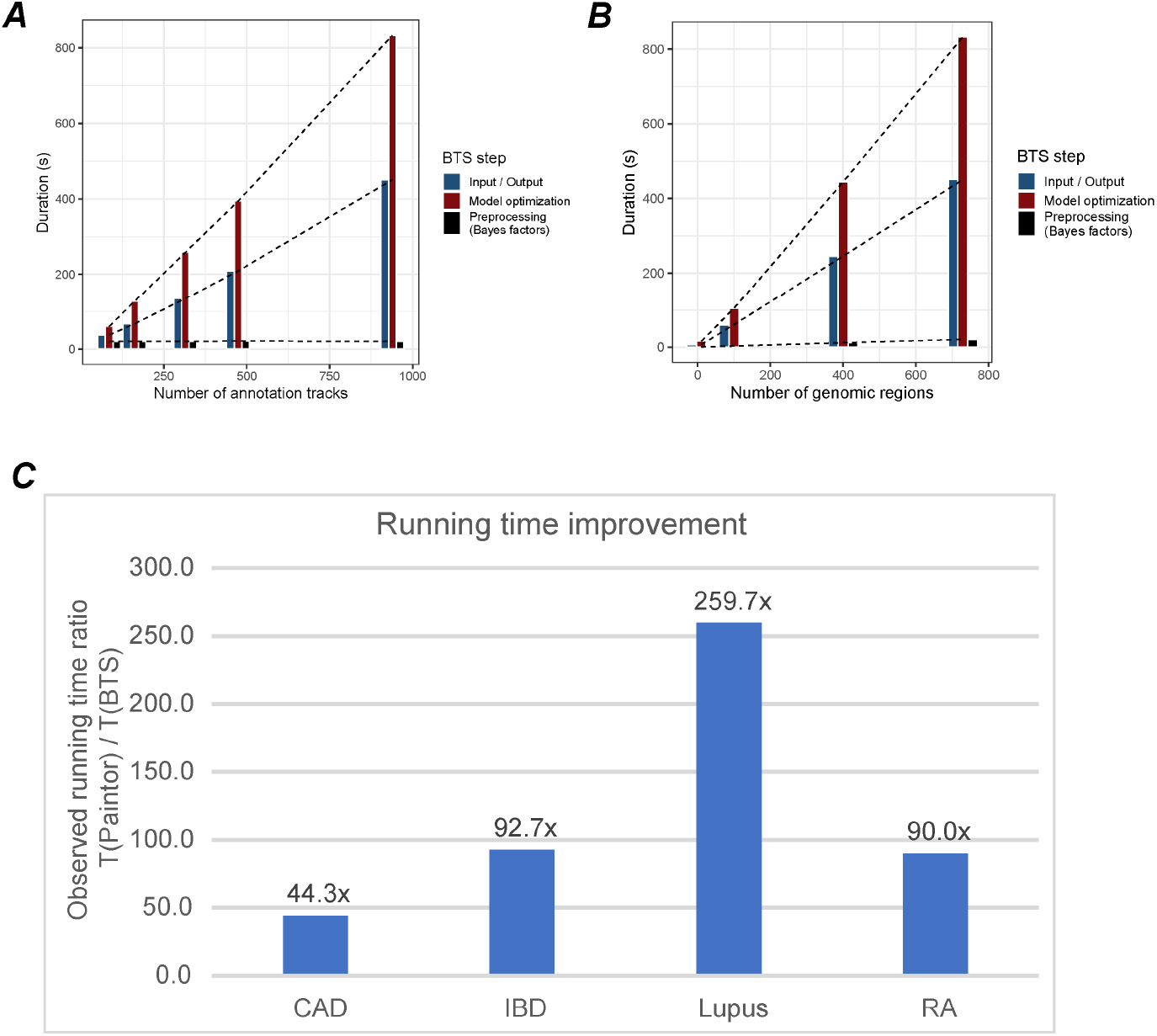
BTS running time. **A**. BTS running time as a function of the number of annotation tracks used for functional evaluation. BTS scales linearly with the number of input annotation tracks. Note the constant running time for pre-computing annotation-independent Bayes factors. **B**. BTS running time as a function of the number of genomic regions. BTS scales linearly with the number of genomic regions of interest. **C**. Comparison of the per-model BTS running time and the running time using the reference implementation (fastPaintor 3.0 (Kichaev et al., 2017)). BTS improves running time by a factor of 44-259x (average 120x) across the four GWAS datasets analyzed.

## Methods

### BTS GWAS summary statistics analysis workflow

We provide an end-to-end functional variant fine-mapping and context-mapping pipeline for analysis of GWAS summary statistics based on user-supplied full GWAS summary statistics as input (**Fig. 1; Supplementary Fig. S1; Supplementary Methods**). The BTS GWAS summary statistics pipeline aims to automate genome-wide post-GWAS functional analysis and provide a systematic and more complete report of all potentially causal variants, genomic loci, and likely functional contexts including cell type and tissue-specific regulatory mechanisms underlying observed GWAS signals.

To do this, input GWAS summary statistics are first preprocessed to resolve reference and non-reference (alternative) alleles and normalize effect sizes to consistently reflect effects of the alternative alleles. Using the normalized GWAS summary statistics, a set of genomic regions for further downstream analyses and all potential candidate variants are then identified. This is achieved through identification of pairwise-independent GWAS signals (*tag* variants) by performing linkage disequilibrium-based pruning (LD r^2^>0.7) of all genome-wide significant (*p* < 5e-8) GWAS variants. Based on the identified tag variants, the candidate set of potentially causal variants is then formed as a set of variants in LD with the tagging variants (LD r^2^>0.7), including the tag variants themselves. LD-based genomic regions for analysis are constructed by defining the LD region for each of the tag variants as a genomic region with the left and right boundaries corresponding to the leftmost and rightmost variants linked with the tag variant. The leftmost and rightmost variants are restricted to be within 1Mbp from the tag variant and have no more than 1000 variants between them and the tag variant. The final set of non-overlapping genomic regions for downstream analyses is obtained by merging any overlapping LD-based regions into larger regions.

For each such identified genomic region, the pipeline will then generate all the information required to fit the BTS model (**Section** “BTS statistical model”) including the pairwise LD-based variant correlation matrix L (*n* x *n*, where *n* is the number of variants in the region), functional annotation matrix A (*n* x N_A_), and a vector of GWAS summary statistics Z (Z-scores) (*n* x 1).

Pairwise LD calculation for all variants located in the locus is conducted based on the reference genotype panel (Byrska-Bishop et al., 2022; Genomes Project et al., 2015). Functional annotation matrix **A** is obtained by querying FILER FG database (Kuksa et al., 2022) for each of the N_A_ genomic annotations and FG data tracks of interest and noting annotation overlaps for each of the variants (**A**_i,j_ will be set to 1 if variant *i* overlaps annotation *j*). The summary of included annotations and detailed list of annotation tracks used are provided in **Supplementary Tables S2**,**S4**. Summary statistics (Z-scores) are extracted from the input GWAS summary statistics after reference and alternative allele resolution and effect normalization to consistently reflect the effect of the non-reference (alternative) allele.

BTS algorithm (**Fig. 1**; **Methods; Section “**BTS algorithm**“**) will then be applied to fit the model and estimate variant and locus posteriors and functional annotation enrichment for each of the target annotations (**Fig. 3**). To find potentially causal variants within each locus, BTS uses variant LD matrix and Z-scores from GWAS summary statistics to pre-compute and store Bayes factors for each possible causal variant configuration in every locus. BTS then uses an EM-based algorithm to iteratively estimate annotation enrichment coefficients and compute annotation-specific causal priors for each of the analyzed variants. These functional annotation-specific priors are then combined with pre-computed configuration Bayes factors to obtain context-specific causal variant posteriors.

Evaluation results reported in **Sections** “Prioritizing regions, variants and their contexts with BTS”, “Cross-trait BTS evaluation” were generated by applying this pipeline to CAD (van der Harst & Verweij, 2018), IBD (Liu et al., 2015), RA (Bentham et al., 2015), and SLE (Stahl et al., 2010) GWAS summary statistics.

### BTS statistical model

To accommodate for correlation between variants (linkage disequilibrium, LD) as well as the tissue and cell-type-specific functions of variants and genomic regions, we adopt the Bayesian probabilistic framework first proposed in (Kichaev et al., 2014). We consider the following information for all *n* variants in a genomic region of interest: 1) a vector **Z** of GWAS Z-scores (standardized regression coefficients; observed), 2) a vector ***A*** of variant tissue and cell type-specific annotations (observed), 3) LD (linkage disequilibrium) correlation matrix **Σ**, and a vector **C** of variant causal status (unobserved; latent binary variable: set to 1 for causal variants, and 0 otherwise), a vector **Λ** of unknown true effect sizes for each variant. To model each genomic region, we use a Bayesian model in which the likelihood of observing **Z** is a multivariate normal parametrized by causal configuration **C**, true effect sizes **Λ**, and LD correlation matrix **Σ**:

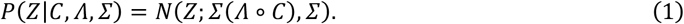

where *Λ* ∘ *C* is an element-wise vector product.

The true effect size **Λ** is also modeled as normal, with mean 0 and diagonal variance, using the scalar *W* as a model parameter:

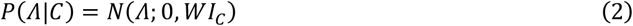

where *WI*_*C*_ is a scaled diagonal matrix, with diagonal elements of *I*_*C*_ set to 0 and 1 according to the causal configuration C.

Integrating out **Λ** gives the following formula for the full likelihood as proved in (Kichaev et al., 2014):

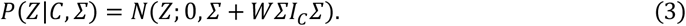

Bayes factor (BF) for a causal variant configuration C in any particular genomic region:

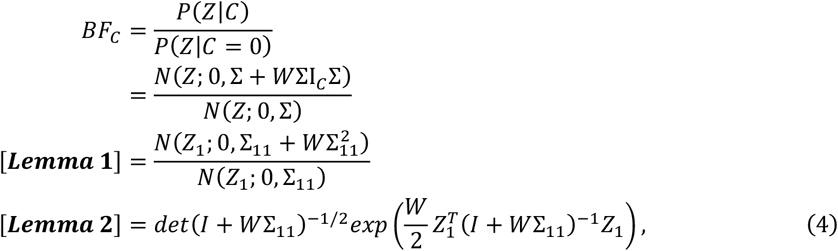

where the first simplification (**Lemma 1)** reduces BF computation from full vectors and matrices to the much smaller 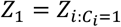, and 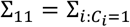 corresponding to the Z-scores and correlation between the causal variants (*C*_*i*_ = 1), and the second simplification (**Lemma 2**) further reduces BF computation to a single matrix inversion of a positive semi-definite matrix (see **Supplementary Methods**).

The prior probability of causality for each variant *i* is modeled as a logistic function of its annotation *A*:

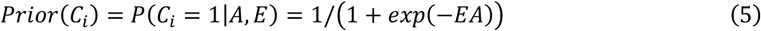

where the annotation effect size coefficient *E* for annotation *A* is estimated genome-wide and is shared by all variants in all regions. Note that for variants overlapping annotation A with positive effect E, their prior probability of causality will be greater than prior probabilities for variants located outside of annotation A. Prior for causal configuration *C*_*j*_ for region *j* is then a product of all *n* individual variant priors *P*(*C*_*i*,*j*_)

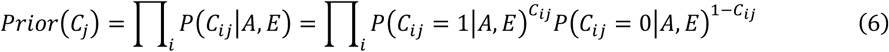

The full data likelihood across all genomic regions *j* is a product of individual region data likelihoods

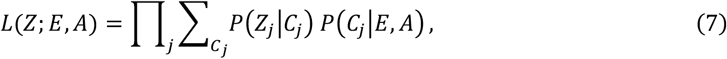

The computational complexity for each region *j* is then consists of prior computation and data likelihood computation (Eq. 3) for every possible causal variant configuration *C, O*(|*C*| × (*L* + *d*^3^)).

To improve computational efficiency, we first note that *P*(*C*_*i*_ = 0|*A, E*) = 1 - *P*(*C*_*i*_ = 1|*A, E*) = 1/(1 + *exp*(*EA*)) and a full variant configuration probability can be computed in *O(d)* time (where *d* is the maximum number of independent causal variants) as an update to the null configuration probability:

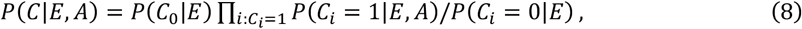

where C_0_ is a null configuration (all variants are non-causal) and the P(C_0_) term is only computed once per locus.

Marginalizing over C, and restricting to configurations that have at most *d* causal variants, we obtain the posterior probability that a variant *i* is causal in a particular genomic region:

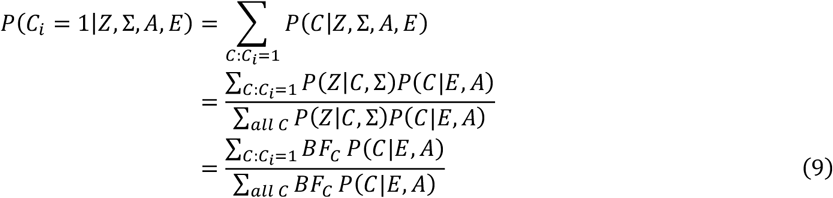

expressed in terms of the Bayes factors *BF*_*C*_ = *P*(*Z*|*C*)/*P*(*Z*|*C*_0_) (Eq. 4) and configuration priors P(C) (Eq. 6).

More conceptually, the variant posterior in (Eq. 9) is a dot product between a vector of annotation-independent Bayes factors *BF*_c_ and a vector of annotation-dependent variant priors *P*(*C*), where each vector is indexed by causal variant configurations C with up to *d* causal variants (i.e. each vector is of 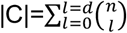 dimensionality, where *n* is the number of variants in the locus). These vectors can be computed independently from each other, as Bayes factors (BF) only depend on GWAS summary statistics and LD matrix (Eq. 4), while the variant priors only depend on annotations and their enrichment coefficients (Eq. 5,6).

Given access to the precomputed Bayes factors for each possible configuration C, the overall complexity of computing variant posteriors (using Eq. 8,9) is then linear *O*(|*C*| × *d*) for any given genome-wide annotation *A*, which is a *O*(*L* + *d*^2^) improvement compared to non-factorized model (Eq. 7) *O*(|*C*| × (*L* + *d*^3^)) with the on-the-fly prior and Bayes factor computation.

The model outputs the variant causal posterior probabilities (Eq. 9) for each of analyzed variants and the estimated annotation effect size coefficients *E*_*A*_ for each tested annotation *A*.

### BTS algorithm

We use an expectation-maximization (EM) algorithm (Kichaev et al., 2014) to fit the statistical model in **Eq. 9**. Intuitively, this is an iterative algorithm which optimizes overall likelihood and takes turns updating the posterior probabilities and enrichment coefficients, until a convergence criterion is reached.

Given multiple annotations, with a possibly complex correlation structure, it is standard practice (Kichaev et al., 2014; Pickrell, 2014) to perform feature selection, by first fitting a separate model for each annotation, and then selecting a few high-ranking annotations for a final model. Therefore, our systematic approach to annotations requires that thousands of models be fitted for all annotation, tissue, and cell types.

Our key observation is that, in (**Eq. 9**), the posterior probability decomposes into a factor which only involves the GWAS data, and one which only involves annotations. Furthermore, the factor involving GWAS data is the same for all iterations of the EM algorithm. Because of this, BTS computes it only once, and distributes it to all models and EM iterations as necessary. Since likelihood computation is the most time-intensive part of fitting the model, the choice to only perform it once is responsible for the bulk of our computational improvements.

This algorithm design choice is a trade-off between speed and memory: to compute the likelihood only once, BTS must store it until all models have been processed. The amount of time and memory spent on likelihood computations is proportional to the number of allowed causal configurations:

- If there are N variants, and any subset of them could be the causal set, then there are 2^N^ causal sets to be enumerated. Due to the nature of exponential growth, this is impractical even for moderately large regions (N>30) and impossible for large ones (N>80).
- BTS implements a common solution to this problem, which is to only consider configurations of size smaller or equal to *d* (Asimit et al., 2019; Kichaev et al., 2014), a user-provided parameter, with default value 2. Then storing the likelihood of such configurations requires O(N^d^) memory for a region with N variants.
- If *d*=2, the necessary memory is equal to that of storing the LD matrix, so BTS gains two orders of magnitude in speed, at the cost of only doubling its memory use.
- For *d*>2, we mitigate the memory use by only storing those likelihoods which are at most t orders of magnitude smaller than the largest, where t is a user-provided parameter, with default value 12. In our experiments with *d*=3,4,5, this optimization reduces memory use by 100-fold, and does not change the final results within the first 5 significant digits. Our experiments suggest that BTS can accommodate values of *d* up to 5 without significant memory issues, while for *d*>5 runtime increases severely.

We obtain further computational improvements by using the matrix inversion lemma (**Lemma 2**) to compute Bayes factors (**Supplementary Methods**) and more efficiently computing variant configuration probabilities (**Eq. 8**). When computing likelihood ratios, BTS needs to evaluate the ratio of multivariate normal densities with different variance matrices. Naively, this involves inverting each variance matrix. The matrix inversion lemma provides an equivalent expression in which a single matrix needs to be inverted and has the following benefits:

- Decreased computation time, since matrix inversion is the most time-consuming part of likelihood computation.
- The covariance matrices are singular in the case of variants in perfect LD. In our formulation, the matrix to be inverted is strictly positive definite, which improves numerical stability and removes the need for regularization.

**Figure 1** summarizes the BTS algorithm:

- A module for computing Bayesian factors and likelihoods.
- A loop which distributes annotations and likelihoods to each model to be fit.
- Aggregation of results, and prioritization of annotations, loci and variants.

BTS GWAS summary statistics pipeline using core BTS algorithm is outlined in **Supplementary Figure S1**.

### Mitigation of LD mismatch

We investigated the effects of LD mismatch, which can occur whenever in-sample LD for the GWAS cohort is unavailable, and a reference genotype panel is used instead. There are existing methods to flag regions with high suspicion of LD mismatch (DENTIST (Chen et al., 2021), SLALOM (Kanai et al., 2022)), but they do not address the problem after identifying it. Moreover, SLALOM only flags a region if the mismatch involves the variant with highest association, which is not completely general.

Our approach is to quantify the spurious effect of LD mismatch on likelihood computations and provide guidance in choosing algorithm parameters so that this effect is minimized. In the supplementary material (see **Supplementary Methods**), we show that, for two variants in perfect LD, with Z-scores a, b, the likelihood of the configuration where both are causal is:

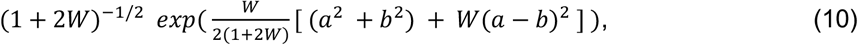

where W is the prior variance from (Eq. 2). Since the variants are perfectly correlated, it should be the case that a=b in the absence of LD mismatch. In practice, we often observe LD=1 but a≠b; one Z-score could be large while the other is close to zero. In this case, the term W(a-b)^2^ in the exponent is the spurious effect which should be minimized. If W is much larger than 1, then the spurious second term can end up dominating the first one. However, if W is much smaller than 1, then the null configuration can end up dominating all others, leading to posterior probabilities close to 0. To balance these requirements, BTS uses W=1.

## Discussion

In this paper we report BTS, a new algorithm that performs joint fine-mapping of variants and context-mapping using genome-wide functional annotations. The algorithm has multiple statistical and algorithmic innovations to allow one to characterize genetic association signals across the genome against thousands of functional assay experiments across different tissue and cell types. The algorithm provides easily interpretable results that highlight important cellular and tissue context of genetic trait associations with genome-wide enrichment and statistical confidence. The BTS algorithm implementation is highly scalable and can allow multiple (up to *d*=5) independent association signals in each locus.

We applied BTS to summary statistics from four GWASs (immune-related and cardiovascular) and compared them against different publicly available functional genomic annotations across >200 tissue and cell types. BTS successfully prioritized relevant tissue and cellular types known to be associated with the disease biology, validating the underlying statistical model and showing the value of our approach to translate genetic findings to biological mechanisms of disease.

BTS has certain limitations. First, currently available functional annotations are still limited in their specificity in cell and tissue types, and this remains to be addressed in the future as the research community continues to generate highly specific functional experiments. Additionally, users can provide their own specialized functional annotations when running BTS. Second, the number of independent causal variants (parameter *d*) per locus needs to be chosen in advance and the memory use is still exponential, which prohibits analysis of high-order genetic interactions. In practice this might not be a serious limitation as the algorithm is still scalable up to *d*=5.

There are several useful usage scenarios for BTS by biologists, geneticists and other researchers. As complex diseases involve multiple cell types and their interactions, the choice of input tracks and genomic features can reflect multiple hypotheses involving suspected or potentially causal tissues, cell types, and disease mechanisms. BTS can then be used to test these hypotheses and select and prioritize tissues, cell types, genomic mechanisms, functional genomic regions, and variants. We note that compared to enrichment-based frameworks such as LDSC (Bulik-Sullivan et al., 2015) and S-LDSC (Finucane et al., 2015) that are commonly used to gain insight into potentially relevant cell types, tissues, BTS not only identifies relevant functional context(s), but also allows to identify and prioritize functional genomic regions for each of these potential contexts and provides fine-mapping/prioritization for variants within each of the analyzed genomic regions and for each of analyzed contexts. While multiple overlapping tracks are typically used to show or prioritize functional regions and individual variants (e.g. in the genome browser), BTS can also output per-annotation or per-locus variant posteriors which can be used to select variants with high causal posteriors across annotations. Overall, BTS provides data-driven discovery and prioritization by combining GWAS signals and FG and annotation data.

There are many directions for future development of this work. First, although BTS ranks region and annotation pairs in its output, it does not rank regions by themselves, nor does the algorithm provide affected target genes (e.g., as in Activity-by-contact (Fulco et al., 2019), Effector Index (Forgetta et al., 2022) models) regulated by the causal variants. Incorporation of expression QTL data or chromatin interaction data could address the second problem, and this idea is explored in many other papers including the INFERNO algorithm we reported previously (INFERNO (Amlie-Wolf et al., 2018), FUMA (Watanabe et al., 2017), SparkINFERNO (Kuksa et al., 2020)). BTS may also be expanded to accommodate for other types of annotations such as experimental functional assays (MPRAs) at gene or transcript level, and predicted functional activity or pathogenicity of variants (e.g., GENO-NET (He et al., 2018), CADD (Rentzsch et al., 2021; Rentzsch et al., 2019); JARVIS (Vitsios et al., 2021); PoPS (Weeks et al., 2023)), but new statistical models need to be developed to integrate these types of data.

An advantage of BTS is its scalability which allows us to explore thousands of genome-wide functional annotations and systematically explore the biological mechanism and cellular and tissue context without bias. One may extend our algorithm to analyze multiple traits and tissues jointly or carry out combinatorial analysis of multiple cell type and tissue functional surveys across many genomic features. Applications of such approach to the growing body of GWAS and WGS-based studies, along with further experimental validations, could lead to significant insights into disease-underlying variants, loci, and molecular mechanisms.

## Supporting information

BTS Supplementary Information

BTS Supplementary Tables

## Data availability

GWAS summary statistics used in this study were obtained from EMBL-EBI GWAS catalog for IBD (https://ftp.ebi.ac.uk/pub/databases/gwas/summary_statistics/GCST003001-GCST004000/GCST003043/IBD_trans_ethnic_association_summ_stats_b37.txt.gz), RA (http://ftp.ebi.ac.uk/pub/databases/gwas/summary_statistics/GCST000001-GCST001000/GCST000679/stahl_2010_20453842_ra_efo0000685_1_gwas.sumstats.tsv.gz), and SLE (http://ftp.ebi.ac.uk/pub/databases/gwas/summary_statistics/GCST003001-GCST004000/GCST003156/bentham_2015_26502338_sle_efo0002690_1_gwas.sumstats.tsv.gz), and from MRC IEU OpenGWAS database for CAD (https://gwas.mrcieu.ac.uk/files/ebi-a-GCST005195/ebi-a-GCST005195.vcf.gz). ENCODE (https://encodeproject.org), Roadmap Epigenomics (http://www.roadmapepigenomics.org/), and EpiMap (https://compbio.mit.edu/epimap/) functional annotations used in BTS evaluations and 1000 Genomes (https://www.internationalgenome.org/) reference genotype panel data for LD computation were obtained from the FILER database (https://lisanwanglab.org/FILER). BTS results on tested GWASs are available from https://doi.org/10.5281/zenodo.14521100. BTS runs on HPC clusters, single servers or cloud-based instances and is available at https://hub.docker.com/r/wanglab/bts (Docker container) and https://bitbucket.org/wanglab-upenn/bts-pipeline (BTS pipeline source code). BTS core R package is also available at https://bitbucket.com/wanglab-upenn/BTS-R.

## Acknowledgments

We thank Wang lab members, Mingyao Li, and Struan Grant for their valuable feedback and comments.

