## Supplementary material for "BTS: scalable Bayesian Tissue Score for prioritizing GWAS variants and their functional contexts across omics data": BTS Supplementary Information

### Supplementary Tables

**Supplementary Table S1**. Comparison of BTS and other methods.

**Supplementary Table S2**. Functional genomic and annotation data used for GWAS evaluation with BTS.

**Supplementary Table S3.** Summary of BTS results for evaluated GWAS datasets

**Supplementary Table S4**. Metadata for all FILER annotation tracks used in BTS analyses.

**Supplementary Table S5**. Prioritized annotation tracks, genomic regions, and variants for (a) CAD, (b) IBD, (c) SLE, (d) RA.

**Supplementary Table S1**. Comparison of BTS and other methods.

| Method | Pipeline Features | | | | Pipeline Outputs | | | |
|---|---|---|---|---|---|---|---|---|
| | **GWAS summary statistics pipeline** | **scalable across many annotation tracks** | **exhaustive causal variant search** | **broad cell type, tissue survey** | **context mapping** | **context-specific variant fine-mapping** | **context-specific locus prioritization** | **References** |
| INFERNO | yes | yes | no | yes, expandable | partial (annotation overlaps) | no | no | (Kuksa et al., 2020) |
| FUMA | yes | web-based | no | limited, non-expandable | partial (annotation overlaps) | no | no | (Watanabe et al., 2017) |
| FINEMAP | no | no | no | no | no | no | no | (Benner et al., 2016) |
| SuSiE | no | no | no | no | no | no | no | (Wang et al., 2020) |
| PAINTOR | no | no | yes | no | only small number of tracks | only small number of tracks | no | (Kichaev et al., 2014) |
| PolyFun | no (only partial) | yes | yes (via PAINTOR) | no | no | yes (via PAINTOR) | no | (Weissbrod et al., 2020) |
| CARMA | no | yes | no | no | no | yes | no | (Yang et al., 2023) |
| BTS | yes | yes | yes | yes, expandable | yes | yes | yes | this paper |

**Supplementary Table S2**. Functional genomic and annotation data used for GWAS evaluation with BTS. All annotation tracks were obtained from FILER (Kuksa et al., 2022). Data source column indicates original data source. See **Supplementary Table S4** for full track information.

| **Genomic feature type** | **Data source** | **#annotation tracks** | **#cell types** | **#tissue categories** | **#annotated genomic regions** |
|---|---|---|---|---|---|
| Enhancers | Roadmap – ChromHMM (Roadmap Epigenomics et al., 2015) | 78 | 65 | 15 | 7,877,498 |
| | EpiMap (Boix et al., 2021) | 316 | 110 | 22 | 38,053,041 |
| | FANTOM5 (Andersson et al., 2014) | 107 | 77 | 28 | 185,397 |
| Open chromatin regions | ENCODE (Consortium, 2012) | 203 | 55 | 19 | 33,463,914 |
| | Roadmap (Roadmap Epigenomics et al., 2015) | 63 | 22 | 7 | 24,506,793 |
| Active histone marks | ENCODE (Consortium, 2012) | 176 | 47 | 20 | 10,956,718 |
| | **TOTAL** | 943 | 194 | 28 | 115,043,361 |

**Supplementary Table S3.** Summary of BTS results for evaluated GWAS datasets

| | | CAD (van der Harst & Verweij, 2018) | IBD (Liu et al., 2015) | SLE (Bentham et al., 2015) | RA (Stahl et al., 2010) |
|---|---|---|---|---|---|
| GWAS | GWAS variants | 7,917,998 | 10,589,089 | 7,910,110 | 2,525,781 |
| Regions of Interest | genome-wide significant variants | 4,298 | 11,967 | 15,980 | 3,835 |
| | tag variants | 389 | 885 | 1,520 | 753 |
| | regions of interest | 167 | 160 | 147 | 58 |
| | variants per region | 148 | 236 | 200 | 91 |
| Prioritization | annotations | 26 | 22 | 11 | 5 |
| | regions | 68 | 52 | 44 | 23 |
| | variants | 84 | 63 | 44 | 27 |
| | variant-annotation pairs | 1,866 | 651 | 97 | 53 |
| | with increased LL | 427 | 372 | 41 | 11 |

**Supplementary Table S4**. Metadata for all FILER annotation tracks used in BTS analyses. See **Supplementary_Tables.xlsx.**

**Supplementary Table S5**. Prioritized annotation tracks, genomic regions, and variants for (a) CAD, (b) IBD, (c) SLE, (d) RA. See **Supplementary_Tables.xlsx.**

### Supplementary Figures

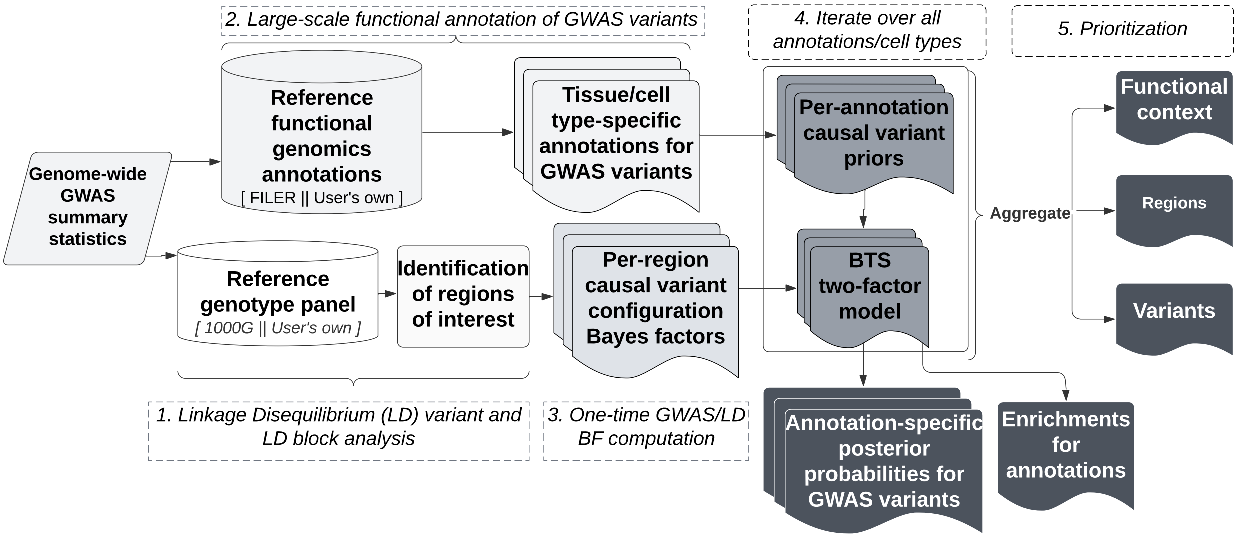


**Supplementary Figure S1**. Overview of BTS GWAS summary statistics analysis pipeline. Input GWAS summary statistics are first analyzed to determine a set of genomic regions for further functional analysis and fine-mapping (Step 1; **Section** "BTS GWAS summary statistics analysis workflow"**; Methods; Supplementary Methods**). Then, for each of identified genomic regions all the corresponding GWAS variants are annotated with genome-wide tissue and cell type-specific annotations of interest (Step 2; Section "BTS GWAS summary statistics analysis workflow"; **Supplementary Methods**). Computed per-locus variant LD correlation matrices, variant GWAS Z-scores, and variant functional annotations (Steps 1,2) are used as input to BTS model (Section "BTS statistical model"). BTS first pre-computes Bayesian Factors for all possible causal variant configurations across all loci (Step 3; **Section** "BTS statistical model") and is then applied iteratively (Step 4; **Section** "BTS GWAS summary statistics analysis workflow") for each of the provided functional annotations. BTS outputs annotation relevance (enrichments), locus-level probabilities, and context-specific causal variant posterior probabilities. The two-factor structure of BTS model (Steps 3, 4; Section "BTS statistical model") allows it to be evaluated very efficiently across thousands of genome-wide annotations (**Fig. 6; Section** "Running time improvement"; **Methods**). Each of the provided annotations, identified genomic regions, and GWAS variants within these regions are then prioritized by BTS based on their estimated enrichments (prior odds), observed increase in causal posterior likelihood, and variant posteriors, respectively (Step 5; **Fig. 3**; **Sections "**Prioritizing regions, variants and their contexts with BTS**", "**Cross-trait BTS evaluation**"**).

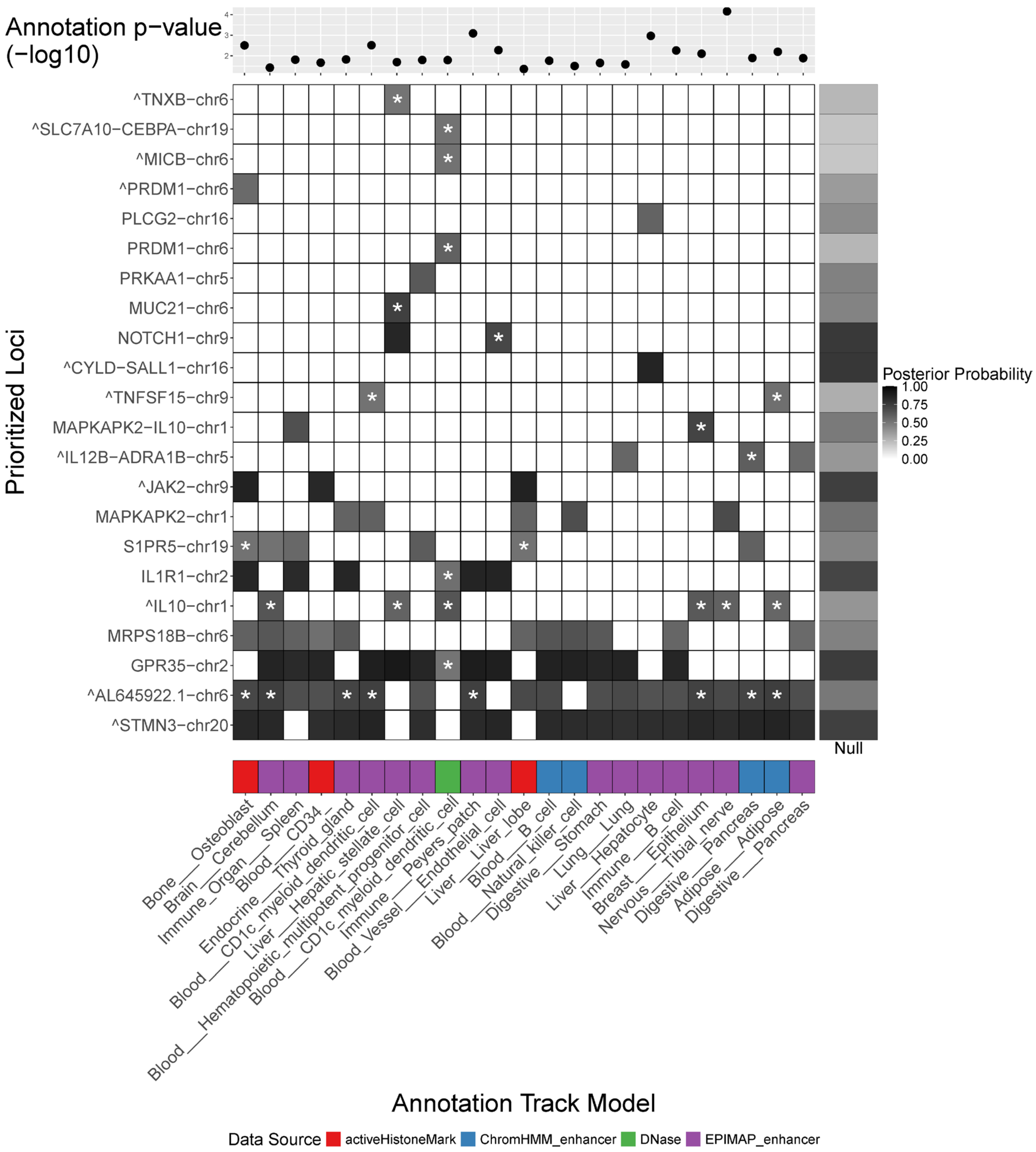


**Supplementary Figure S2**. BTS prioritizes digestive, blood, and immune tissues and annotations based on IBD GWAS summary statistics. Shown are prioritized genomic regions (Y axis) and their functional contexts (X axis). For each region and context shown is the top variant posterior (shades of gray) with the star (*) indicating posterior increase of at least .2 in that context compared to the null model without annotation (right).

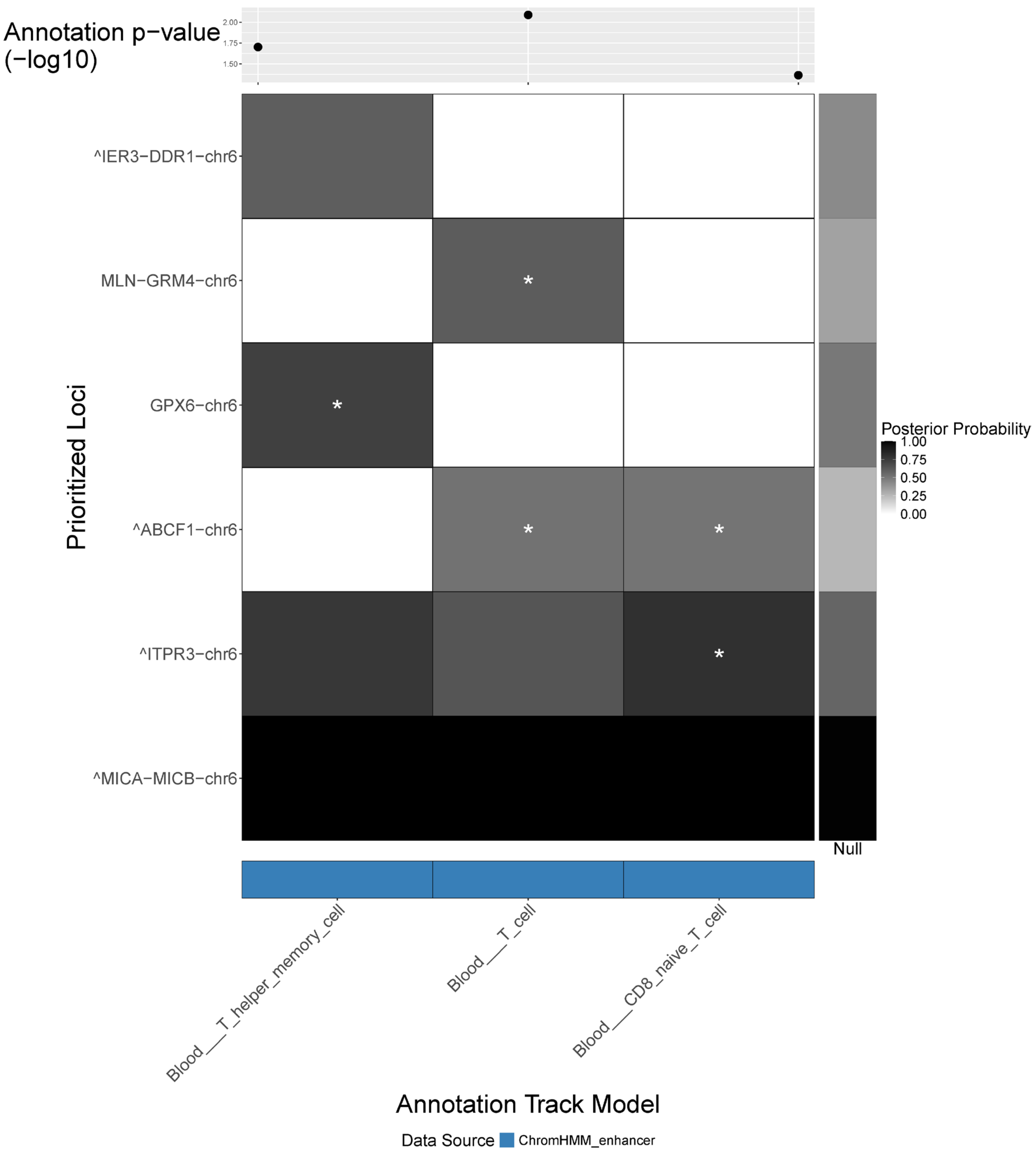


**Supplementary Figure S3**. BTS prioritizes T cells and ChromHMM enhancer annotations for RA. Shown are prioritized genomic regions (Y axis) and their functional contexts (X axis). For each region and context shown is the top variant posterior (shades of gray) with the star (*) indicating posterior increase of at least .2 in that context compared to the null model without annotation (right).

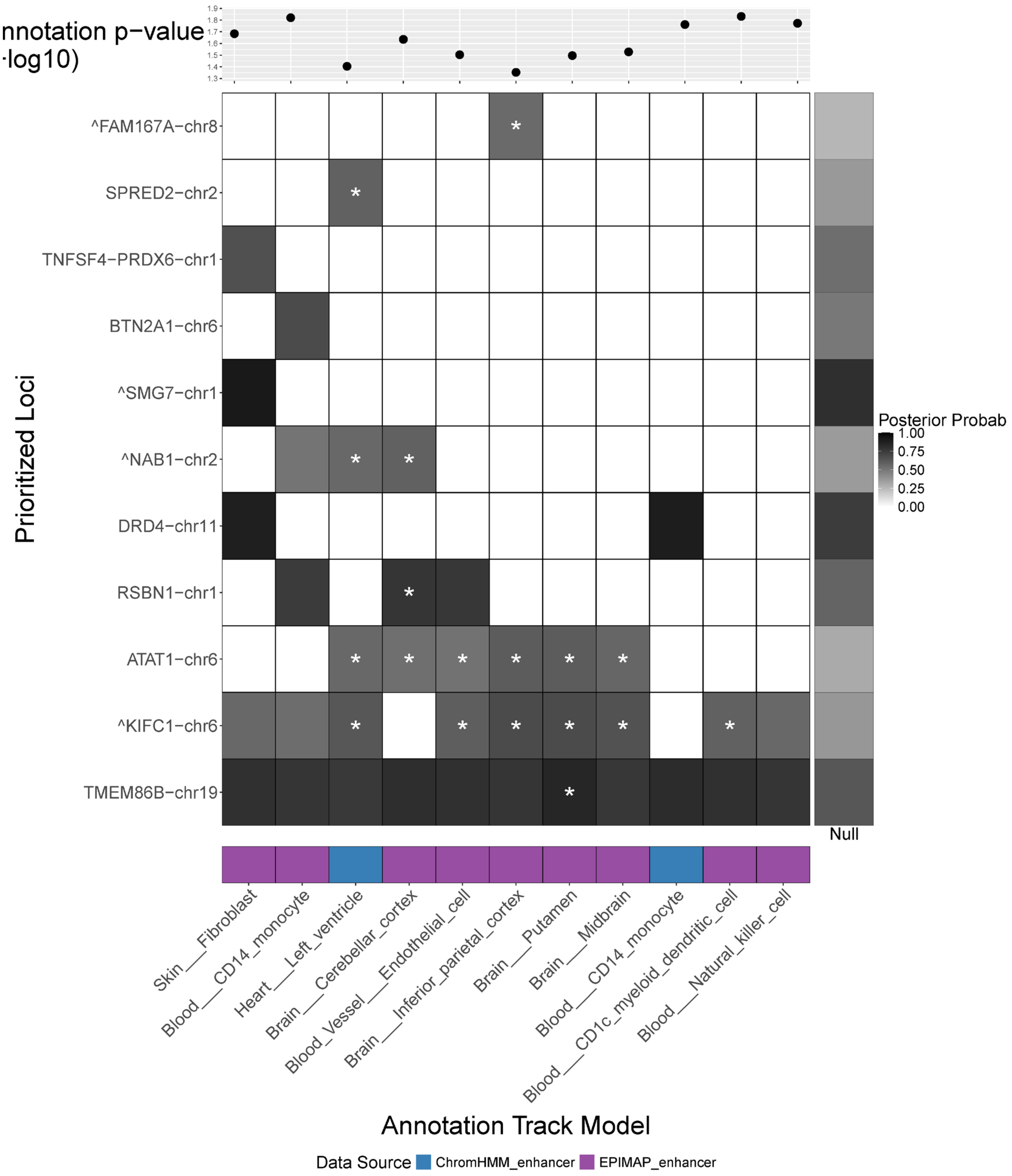


**Supplementary Figure S4**. BTS prioritizes heart left ventricle, monocyte, and brain tissues and annotations based on SLE GWAS summary statistics. Shown are prioritized genomic regions (Y axis) and their functional contexts (X axis). For each region and context shown is the top variant posterior (shades of gray) with the star (*) indicating posterior increase of at least .2 in that context compared to the null model without annotation (right).

Immune-related category prioritized by BTS.

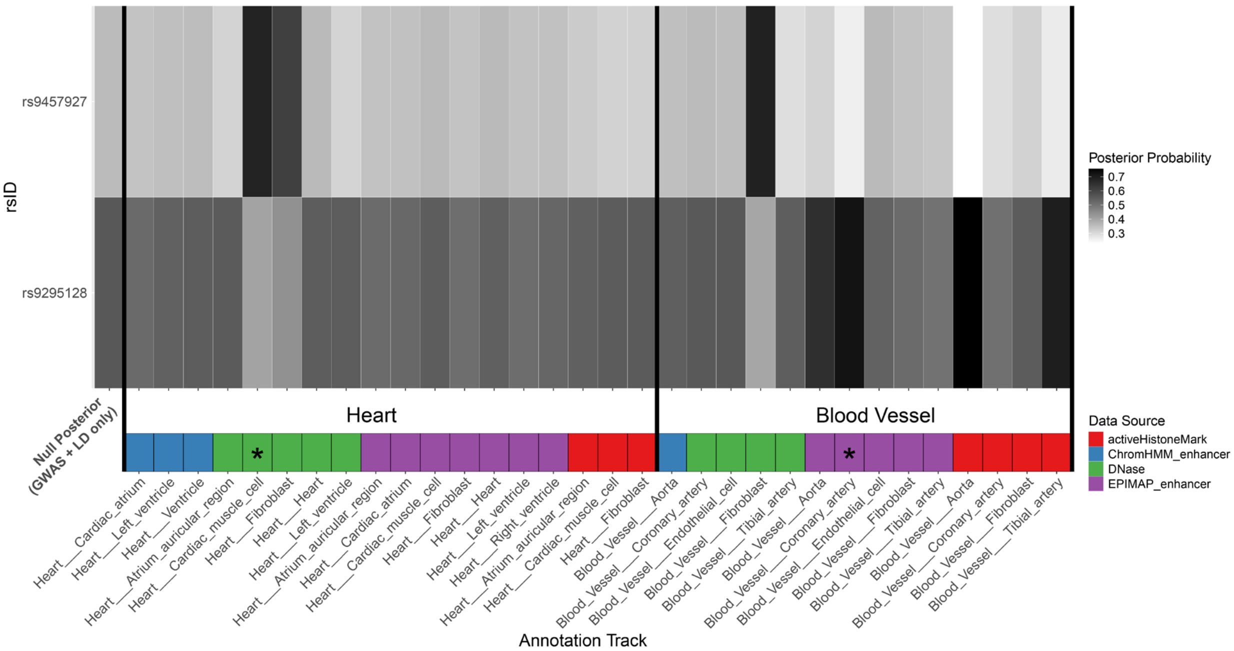


**Supplementary Figure S5**. Differential cell-type and tissue-specific credible set variant prioritization for SLCA22A region. Shown are variant posteriors across heart and blood vessel annotations. Also shown are the baseline variant posteriors (left) for model without any annotations (estimated using GWAS Z-scores and LD information only). Variants rs9295128 and rs9457927 are prioritized in different cell and genomic feature contexts including coronary artery (EpiMap enhancer) and cardiac muscle cell open chromatin (DNase) contexts, respectively. Annotation tracks displayed in **Figure 4** are marked with a star (*).

### Supplementary Methods

#### 1 BTS GWAS summary statistics analysis workflow

##### Input

1. Genome-wide GWAS summary statistics dataset $G$ ($Z$-scores)
2. Reference genotype panel $R$ for LD estimation (e.g., 1000 Genomes EUR super-population) or pre-computed genome-wide pairwise LD scores
3. Reference variant information $V$ [e.g, dbSNP]
4. Reference functional genomic and annotation tracks data collection $D = \{D_1, \ldots, D_K\}$ and corresponding metadata $MD = \{MD_1, \ldots, MD_K\}$ [e.g., FILER]
5. Genomic regions of interest (GWAS loci) $I = \{I_1, \ldots, I_N\}$, where each $I_j = [chr_j, left_j, right_j]$ is a genomic region to be analyzed and fine-mapped. These genomic regions of interest can be either user-supplied or inferred from GWAS summary statistics $G$ and $R$ by GWAS genome-wide signficant variant LD pruning/LD expansion (see **Methods**).

##### BTS GWAS summary statistics analysis algorithm

1. GWAS1. From $G$ and reference variant information database $V$, infer reference and non-reference alleles for each reported variant/association and compute non-reference-allele adjusted summary statistics $G_{std}$ and reference/non-reference variant representations [*Giggle, dbSNP*]
2. GWAS2. Using $G_{std}$ and $R$, obtain regions of interest by first LD pruning (to obtain a set of pairwise independent variants) and then LD-expanding genome-wide significant ($p < 5e-8$) GWAS variants from $G_{std}$ to obtain a set of LD blocks. Merge any overlapping LD blocks to obtain a final set of non-overlapping regions of interest $L_j$, $j = 1 \ldots N$ for further analysis.
3. GWAS3. From $G_{std}$, obtain Z-scores for each locus $j$, $Z_j = \frac{\beta_j^{non-ref}}{se_j}$, $[|L_j| \times 1]$, where $|L_j|$ is the number of variants in locus $j$.
4. LD. Compute pairwise LD matrix $LD_j$, $[|L_j| \times |L_j|]$, for every locus $j$ using $R$ [or extract $LD_j$ from the pre-computed LD scores] [*Giggle, plink*]
5. FG. For each locus $j$, compute variant annotation matrices $A^j_{i,k} = 1, [|L_j| \times K]$ if the $i$th SNP in locus $j$ overlaps $k$th annotation $D_k, k = 1 \ldots K$. Note: annotations maybe selected based on the annotation metadata $MD$. [*FILER*]
6. BTS1. Pre-compute probabilities for all possible causal configuration $C_j$, $[\sum_{i=0}^{i=d} \binom{|L_j|}{i} \times 1]$ for each locus $j$ using $LD_j$ and $Z_j$ [ Bayesian factors $BF_c, c = 1, \ldots, |C_j|$; `compute_configs(LD,Z)`]
7. BTS2. Run EM using the pre-computed configuration probabilities $BF_{C_j}$ and annotation matrix $A^j$, $j = 1, \ldots, L$ [`run_em(P(C), A)`]
   - estimate annotation enrichment/effect sizes $\gamma_k$ for every genomic annotation track $D_k$, $k = 1, \ldots, K$
   - compute variant posteriors $P^j(C_i)$ for every variant $i = 1, \ldots, |L_j|$ and locus $j = 1, \ldots, N$
8. BTS3. Summarize each of the $K + 1$ estimated models across $L$ loci [`get_model_summary`]

- O1. per-annotation variant causal posteriors $P^j_{i,k}$, $j = 1, \ldots, L$, $i = 1, \ldots, |L_j|$,$k = 1, \ldots, K$
- O2. (optimized) total data likelihoods $LL_k$ for each model $k = 1, \ldots, K$
- O3. annotation relevance / enrichments $E_k, k = 1, \ldots, K$
- O4. annotation-specific locus likelihoods $ll^{locus}_j$ and significance scores $p^{locus}_{j,k}$, $j = 1, \ldots, N$, $k = 1, \ldots, K$

**Main data objects**: Loci $L = \{L_1, ..., L_L\}$, SNPs $S = \{S_1, ..., S_L\}$, summary statistics $Z = \{Z_1, ..., Z_L\}$, locus LD matrices $LD = \{LD_1, ..., LD_L\}$ reference annotation data $D = \{D_1, ..., D_K\}$, metadata $MD = \{MD_1, \ldots, MD_K\}$, variant/loci annotation matrices $\{A^j_{i,k}\}$, $j = 1, \ldots, L$, $i = 1, \ldots, L_j$, $k = 1, \ldots, K$

### 2 Causal variant configuration modeling

Let $Z$ be a vector of GWAS $Z$-scores (standardized regression coefficients), $\Lambda$ a vector of true effect sizes (unknown), $\Sigma$ the correlation (LD) matrix, and $C$ a binary vector of causal variant status. We want to estimate the probability of each causal configuration $C$, conditional on the observed data and the model parameters:

$$P(C|Z, \Lambda, \Sigma). \tag{1}$$

We use Bayes' theorem:

$$P(C|Z, \Lambda, \Sigma) = \frac{P(Z|C, \Lambda, \Sigma) \cdot P(C)}{P(Z|\Lambda, \Sigma)}. \tag{2}$$

On the right hand-side, $P(Z|C, \Lambda, \Sigma)$ is the likelihood of observing the data $Z$ under configuration $C$, $P(C)$ is the prior of the configuration (which will depend on functional annotations of the variants). The denominator is the total data likelihood under the model; to compute it is difficult and unnecessary, because it suffices to consider probability ratios, relative to the null configuration:

$$\frac{P(C|Z, \Lambda, \Sigma)}{P(C = 0|Z, \Lambda, \Sigma)} = \frac{P(Z|C, \Lambda, \Sigma) \cdot P(C)}{P(Z|C = 0, \Lambda, \Sigma) \cdot P(C = 0)} = BF(C, Z, \Sigma)\frac{P(C)}{P(C = 0)}. \tag{3}$$

The Bayes factor is, by definition, the ratio of likelihoods. It will turn out to have a dominant contribution in the posterior probabilities, while the priors act as fine-tuning or tie-breakers. This accurately models our expectation that GWAS summary statistics should carry more weight than functional annotations: if a variant has negligible association, is should never be prioritized.

### 3 Modeling and computing configuration Bayes factors

We model the data likelihood for summary statistics as a multivariate normal:

$$P(Z|C, \Lambda, \Sigma) = \mathcal{N}(Z; \Sigma \cdot \Lambda \circ C, \Sigma). \tag{4}$$

We also model the effect sizes as normal, with mean 0 and diagonal variance, using the scalar $W$ as a model parameter:

$$\Lambda \sim \mathcal{N}(0, W \cdot I). \tag{5}$$

Combining 4 and 5 gives a multivariate normal with mean 0 and variance $\Sigma + Var(\Sigma \cdot \Lambda \circ C)$. Computing the second term gives:

$$P(Z|C, \Sigma) = \mathcal{N}(Z; 0, \Sigma + W \cdot \Sigma C \Sigma). \tag{6}$$

For the null configuration, $C = 0$, and $P(Z|C, \Sigma) = \mathcal{N}(Z; 0, \Sigma)$.

Permuting labels if necessary, we can assume that causal configuration $C = (C_1|C_0)$, where $C_1$ consists of ones (the causal variants) and $C_0$ consists of zeros. Let $Z = (Z_1|Z_0)$ be the decomposition of $Z$ corresponding to causal and non-causal blocks, and similarly for $\Sigma$:

$$\Sigma = \left( \begin{array}{c|c} \Sigma_{11} & \Sigma_{10} \\ \hline \Sigma_{01} & \Sigma_{00} \end{array} \right). \tag{7}$$

### 4 Computing Bayes factors efficiently

Efficient computatio of Bayes factors in BTS is achieved using the following two lemmas/observations.

**Lemma 1** allows to speed-up Bayes factor computation by ignoring all the blocks involving non-causal variants, and work with much smaller arrays ([1], [2]).

**Lemma 1**

$$\frac{\mathcal{N}(Z;0,\Sigma+W\cdot\Sigma C\Sigma)}{\mathcal{N}(Z;0,\Sigma)}=\frac{\mathcal{N}(Z_1;0,\Sigma_{11}+W\cdot\Sigma_{11}^2)}{\mathcal{N}(Z_1;0,\Sigma_{11})}. \tag{8}$$

Our own observation is that this can be further simplified (**Lemma 2**).

**Lemma 2**

$$\frac{\mathcal{N}(Z_1;0,\Sigma_{11}+W\cdot\Sigma_{11}^2)}{\mathcal{N}(Z_1;0,\Sigma_{11})}=\det(I+W\Sigma_{11})^{-1/2}\exp\left(\frac{W}{2}Z_1^T(I+W\Sigma_{11})^{-1}Z_1\right). \tag{9}$$

The advantages of this formulation are:

- it computes a single matrix inverse instead of two, decreasing computation time;
- the matrix being inverted, $I+W\Sigma_{11}$, is always strictly positive definite, even when $\Sigma_{11}$ is singular. This helps with numerical precision, and avoids the need for any regularization of $\Sigma_{11}$.

To prove Lemma 2, start from the left-hand side, writing out the multivariate normal densities:

$$\begin{aligned}&\frac{\det\left(2\pi(\Sigma_{11}+W\Sigma_{11}^2)\right)^{-1/2}\exp\left(\frac{1}{2}Z_1^T(\Sigma_{11}+W\Sigma_{11}^2)^{-1}Z_1\right)}{\det(2\pi\Sigma_{11})^{-1/2}\exp\left(\frac{1}{2}Z_1^T(\Sigma_{11})^{-1}Z_1\right)}\\&=\det\left(\Sigma_{11}^{-1}(\Sigma_{11}+W\Sigma_{11}^2)\right)^{-1/2}\exp\left(\frac{1}{2}Z_1^T\left(\Sigma_{11}^{-1}-(\Sigma_{11}+W\Sigma_{11}^2)^{-1}\right)Z_1\right)\end{aligned} \tag{10}$$

It remains to show that $\Sigma_{11}^{-1}-(\Sigma_{11}+W\Sigma_{11}^2)^{-1}=W(I+W\Sigma_{11})^{-1}$. This is an immediate consequence of the "matrix inversion lemma":

$$A^{-1}+B^{-1}=A^{-1}(A+B)B^{-1}, \tag{11}$$

using $A=\Sigma_{11}$ and $B=-(\Sigma_{11}+W\Sigma_{11}^2)$.

The easiest way to prove Lemma 1 is to apply our strategy from Lemma 2 to both sides.

### 5 GWAS/LD mismatch

For simplicity, assume that there are only two variants and denote:

$$Z=\begin{bmatrix}a\\b\end{bmatrix},\qquad\Sigma=\begin{bmatrix}1&r\\r&1\end{bmatrix}. \tag{12}$$

Let $BF_a$, $BF_b$, $BF_{ab}$ be the Bayes factors for configurations where the first variant, the second variant or both are causal, respectively. In configurations with a single causal variant, the causal block $\Sigma_{11}=\begin{bmatrix}1\end{bmatrix}$; in particular, the Bayes factor is independent of the correlation $r$. According to Lemma 2:

$$\begin{aligned}BF_a&=(1+W)^{-1/2}\exp\left(\frac{W}{2(1+W)}a^2\right),\\BF_b&=(1+W)^{-1/2}\exp\left(\frac{W}{2(1+W)}b^2\right).\end{aligned} \tag{13}$$

(As a side note, replacing $W$ by $W/V$, where $V$ is the variance in the estimate of the regression coefficient, gives the Bayes factors from coloc. So these models are closely related.)

For the configuration with two causal variants, the causal block $\Sigma_{11} = \Sigma$, so:

$$\det(I+W\Sigma_{11}) = (1+W)^2 - W^2 r^2, \qquad (I+W\Sigma_{11})^{-1} = \frac{1}{(1+W)^2 - W^2 r^2} \begin{bmatrix} 1+W & -Wr \\ -Wr & 1+W \end{bmatrix}. \tag{14}$$

$$BF_{ab} = \left((1+W)^2 - W^2 r^2\right)^{-1/2} \exp\left(\frac{W}{2((1+W)^2 - W^2 r^2)}\left[(1+W)(a^2+b^2) - 2Wrab\right]\right) \tag{15}$$

**In the case r=0**, 15 reduces to:

$$BF_{ab}^{r=0} = (1+W)^{-1} \exp\left(\frac{W}{2(1+W)}(a^2+b^2)\right). \tag{16}$$

So:

$$\frac{BF_{ab}^{r=0}}{BF_a} = (1+W)^{-1/2} \exp\left(\frac{W}{2(1+W)} b^2\right). \tag{17}$$

- In other words, when the two variants are uncorrelated, the likelihood ratio of $C_{ab}$ relative to $C_a$ increases exponentially with $b$. This makes sense, because $b$ is bringing new information that cannot be explained by $a$ alone.
- To have significant posterior probability mass on $C_{ab}$, the exponential factor must compete with $(1+W)^{-1/2}$, as well as with the prior, which gets smaller with the inclusion of more causal variants.
- For this to work well in practice, $W$ must not be too small; say $W > 1$.

**In the case r=1**, 15 reduces to:

$$BF_{ab}^{r=1} = (1+2W)^{-1/2} \exp\left(\frac{W}{2(1+2W)}\left[(a^2+b^2) + W(a-b)^2\right]\right), \tag{18}$$

so:

$$\frac{BF_{ab}^{r=1}}{BF_a} = \left(\frac{1+2W}{1+W}\right)^{-1/2} \exp\left(\frac{W}{2(1+2W)}\left[b^2 - \frac{W}{1+W}a^2 + W(a-b)^2\right]\right). \tag{19}$$

Since $r = 1$, we would expect that $b \simeq a$, in which case $W(a-b)^2 \simeq 0$. If $W$ is large enough, then $b^2 - \frac{W}{1+W}a^2$ is also small, and the exponent does not grow too fast. In this case, $C_a$ and $C_b$ end up with more probability mass than $C_{ab}$: the model correctly infers that either variant can be causal, but both are unlikely.

However, if LD is obtained from a reference panel which does not match the GWAS population well, it is possible to have $r = 1$ and $b \not\simeq a$. In this case, as $W(a-b)^2$ becomes large, $BF_{ab}^{r=1}$ dominates not only $BF_a$ and $BF_b$, but also the Bayes factors for variants other than the two considered here. In practice, we even see examples where $r = 1$, but $a$ and $b$ have different signs. In these cases, most of the probability mass ends up on the configuration $C_{ab}$, at the expense of other variants which have stronger GWAS association than either a or b!

In practice, we can mitigate this danger by not letting $W$ be too large, thus controlling the term $W(a-b)^2$; say $W < 20$. Together with the earlier requirement, we find that a reasonable range is $1 < W < 20$.

#### Methods References

[1] Gleb Kichaev, Wen-Yun Yang, Sara Lindstrom, Farhad Hormozdiari, Eleazar Eskin, Alkes L. Price, Peter Kraft, and Bogdan Pasaniuc. Integrating functional data to prioritize causal variants in statistical fine-mapping studies. *PLOS Genetics*, 10(10):1–16, 10 2014.

[2] Farhad Hormozdiari, Martijn van de Bunt, Ayellet V. Segrè, Xiao Li, Jong Wha J. Joo, Michael Bilow, Jae Hoon Sul, Sriram Sankararaman, Bogdan Pasaniuc, and Eleazar Eskin. Colocalization of gwas and eqtl signals detects target genes. *The American Journal of Human Genetics*, 99(6):1245–1260, 2016.
